# Epstein-Barr virus-encoded BART9 and BART15 miRNAs are elevated in exosomes of cerebrospinal fluid from relapsing-remitting multiple sclerosis patients

**DOI:** 10.1101/2023.11.21.568021

**Authors:** Mina Mohammadinasr, Soheila Montazersaheb, Vahid Hosseini, Houman Kahroba, Mahnaz Talebi, Ommoleila Molavi, Hormoz Ayromlou, Mohammad Saeid Hejazi

## Abstract

Epstein-Barr virus (EBV) infection is approved as the main environmental trigger of multiple sclerosis (MS). In this path, we quantified EBV-encoded BART9-3p and BART15 miRNAs in exosomes of cerebrospinal fluid (CSF) of untreated relapsing-remitting MS (RRMS) patients in comparison with the control group. Interestingly, patients displayed significant upregulation of BART9-3p (18.4-fold) and BART15 (3.1-fold) expression in CSF exosomes. Moreover, the expression levels of miR-21-5p and miR-146a-5p were found to be significantly elevated in the CSF samples obtained from the patient group in comparison to those obtained from the HC group. The levels of Interferon gamma (IFN-γ), interleukin-1β (IL-1β), interleukin- 6 (IL-6), interleukin-17 (IL-17), interleukin-23 (IL-23), transforming growth factor beta (TGF- β), and tumor necrosis factor alpha (TNF-α) were observed to be significantly elevated in the serum and CSF exosomes of the patients. The highest increase was observed in TGF-β (8.5- fold), followed by IL-23 (3.9-fold) in CSF exosomes. These findings are in agreement with the association between EBV infection and inflammatory cytokines induction. Furthermore, the ratios of TGF-β:TNF-α and TGF-β:IFN-γ attained values of 4 to 16.4 and 1.3 to 3.6, respectively, in the CSF exosomes of the patients, in comparison to those of the control group. Remarkable stimulation of EBV BART9-3p, BART15 miRNAs, and inflammatory cytokines expression in CSF exosomes confers a substantial link between EBV in MS onset and also the infection-to-MS transition.

## 1. Introduction

Epstein-Barr virus (EBV) belongs to the herpes virus family that infects above 95% of adults worldwide. EBV infection often has not any symptoms and the virus establishes a lifelong latent infection. EBV is associated with many diseases, including Burkitt’s lymphoma, nasopharyngeal carcinoma, [1–4], and autoimmune diseases such as systemic lupus erythematosus, Sjogren’s syndrome, rheumatoid arthritis, and multiple sclerosis [5, 6]. EBV expresses 44 mature microRNAs (miRNAs) derived from 25 miRNA precursors encoded by two primary transcripts (BHRF1 and BART). The binding of miRNAs to 3′-UTR of mRNA lead to inhibit of target genes, so they demonstrate an important role in gene expression regulation [2, 7]. There are two mechanisms in the pathogenesis of MS by EBV miRNAs: (1) targeting MS genes by EBV miRNAs during the transformation of B-cell and (2) dysregulation of host cell miRNAs via EBV infection. Thus, Deregulated host miRNAs alter B cell function. The miRNA expression profile of B cells before and after EBV infection found that EBV miRNAs and host B cell miRNAs were dysregulated after EBV infection and involved in MS pathogenesis through their interaction with MS risk loci [8]. Another study identified the expression of EBV miRNAs in MS and explored potential targets for EBV miRNAs [9]. EBV miRNAs, including BART miRNAs, have essential functions in cancer growth, tumor invasion, and host immune surveillance [10]. For instance, miR-BART15 can attach itself to the 3’-UTR of NLRP3 leading to the downregulation of its downstream cytokines and ultimately reducing inflammation responses [11]. The observation that secretory exosomes are rich in small RNAs, including miRNAs with 19-22 nucleotides, compared to cellular RNAs, and that high levels of EBV-BART miRNAs are loaded in exosomes obtained from EBV-infected cells [12], highlights the of interferon-gamma (IFN-γ), interleukin-1β (IL-1β), interleukin-6 (IL-6), interleukin-17 (IL-17), interleukin-23 (IL-23), transforming growth factor beta (TGF-β) and tumor necrosis factor-alpha (TNF-α) in CSF and serum exosomes were quantified. Since the evidence shows that EBV could induce the secretion of these inflammatory cytokines, they were chosen for this research [24–29].

## 2. Materials and methods

### 2.1. Participants

All research participants attended the Neurosciences Research Center (NRC) located at Imam Reza Hospital, a general hospital of Tabriz University of Medical Sciences (TUOMS), Iran. Written informed consent was obtained from all of the participants. The current investigation was approved by the Ethics Committee of TUOMS (IR.TBZMED.REC.1402.348). All procedures outlined in the study are performed according to ethical standards as they are commonly practiced in clinical settings. MS diagnosis was based on the revised McDonald criteria [30], and the clinical phenotypes were defined by Lublin et al. to differentiate RRMs from PPMS and SPMS [31].

### 2.2. Sample collection and preparation

As previously reported [32], One ml of serum was obtained from each patient at the Plasma Clinical Diagnostic Laboratory located in Tabriz, Iran. Similarly, one ml of serum specimen was collected from each of the thirty individuals who were subjected to routine medical examinations at Imam Reza Hospital and diagnosed as normal subjects, termed healthy controls (HCs). Likewise, CSF samples were obtained from each patient with RRMS from the Plasma laboratory. CSF samples were collected from healthy individuals with suspected meningitis infection and trauma, who were referred to Imam Reza’s Central Laboratory and Sina Hospital’s Central Laboratory from 2018 to 2021, and then were found normal and healthy in terms of meningitis infection. To this end, we acquired residual serum and CSF samples from patients diagnosed with RRMS, without the need for extra sampling of this study. The RRMS and HC groups were matched for age, sex, and ethnicity.

### 2.3. Extraction and characterization of exosomes

Exosomes were isolated from serum and CSF specimens using differential centrifugation as described by Mohammadinasr et al. [32]. The isolated exosomes were subsequently resuspended in a solution of 1x phosphate-buffered saline (PBS), pooled together, and stored at a temperature of -80 °C for subsequent analysis [33].

### 2.4. Reverse transcription and qPCR

The extracted exosomal RNA samples were reverse transcribed to cDNA using a specific stem-loop RT primer. In brief, a total volume of 20 μL was prepared with RT buffer (5x), 200 U/μl reverse transcriptase (MMLV), 10 mM dNTP mixture, 40 U/μl RNase inhibitor, and DEPC-treated water. A specific stem-loop reverse transcriptase (RT)-PCR primer was designed for each primer using the sRNAPrimerDB server (http://www.srnaprimerdb.com) as depicted in table 1. The miRNA sequences were obtained from www.mirbase.org. Primers specificity was determined using Oligo7 software (version: 7.60). The program utilized for the cDNA synthesis was implemented by the following protocol: 15 minutes at16°C, 20 minutes at 37°C, 20 minutes at 42°C, and 5 minutes at 95°C.

**Table 1.**
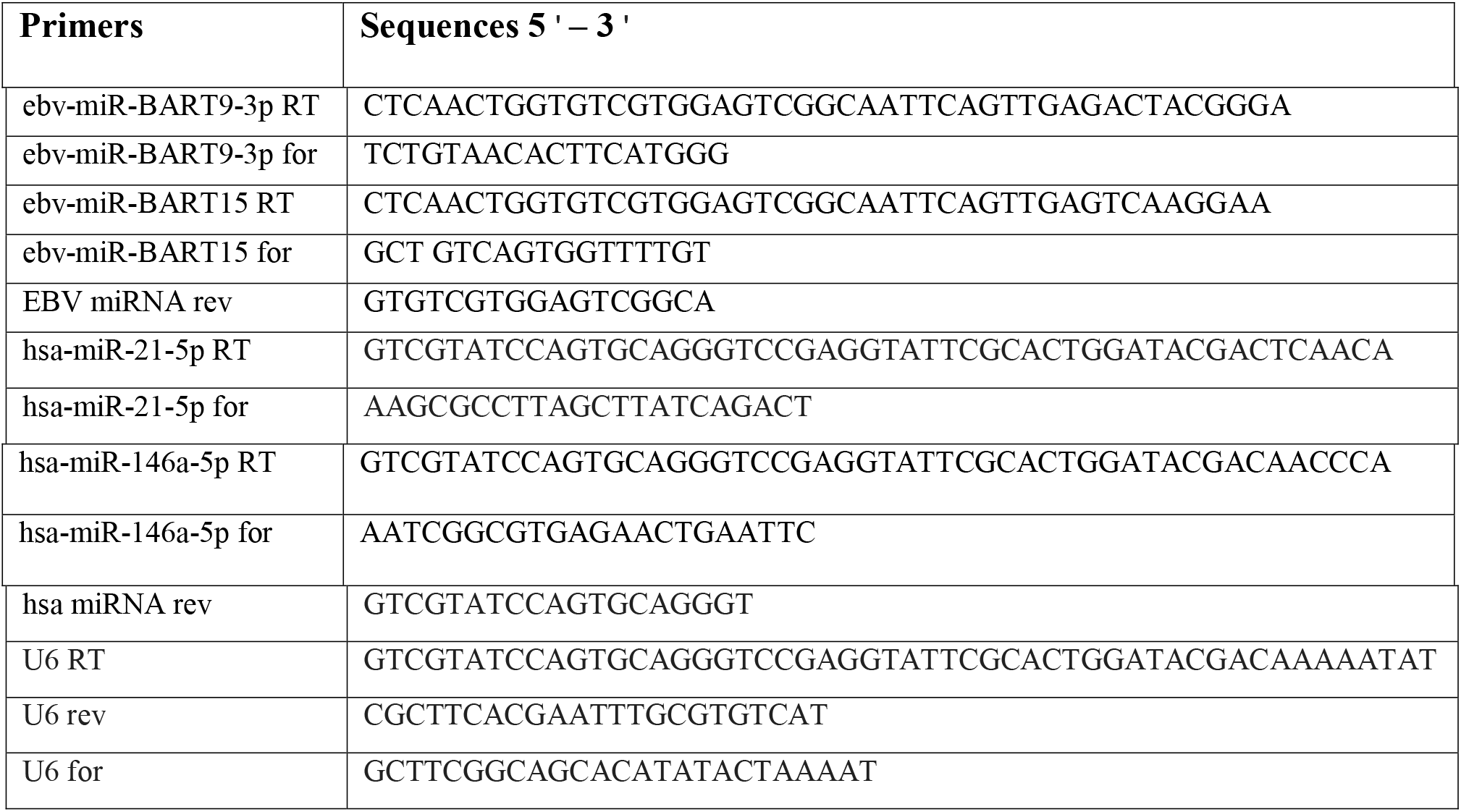
The sequences of the used primers in this study.

Two miRNA primers were used to evaluate the miRNA expression levels using the Corbett Rotor-GeneTM 6000HRM thermocycler. The amplification profile of quantitative Real-Time PCR (qPCR) was as follows: 15 minutes at 95°C for initial denaturation followed by cycles of denaturation at 95°C for 10 seconds and annealing/stretching at 58-62°C (based on the used primer) for 60 seconds. All qPCR was performed in a total volume of 10 μL composed of 5 μL of SYBR Green Master Mix (RealQ Plus 2x Master Mix Green, Ampliqon, Denmark), 1 μL of cDNA, 1 μL of Primer Pair Mix (5 pmol/μl per primer) and 3 µl of distilled water. To quantify the expression of genes, U6 miRNA was employed as an endogenous housekeeping gene for normalizing miRNA expression. Each reaction was conducted a minimum of three times to ensure accuracy. The relative expression levels of miRNA were assessed utilizing the formula _2_−ΔΔCT.

### 2.5. Cytokines assay

To evaluate the levels of IFN-γ, IL-1β, IL-6, IL-17, IL-23, TGF-β, and TNF-α in the exosomes of CSF and serum, we used enzyme-linked immunosorbent assay (ELISA) according to the human DuoSet ® ELISA kits (Catalog Numbers # IFN-γ: DY285, 15.6-1000 pg/mL; IL- 1β: DY201, 3.9-250 pg/mL; IL-6: DY206, 9.4-600 pg/mL; IL-17: DY317, 15.6-1,000 pg/mL; IL-23: DY1290, 125-8000 pg/mL; TGF-β: DY240, 31.2-2000 pg/mL; TNF-α: DY210, 15.6-1000 pg/mL; R&D Systems, USA). The absorbance was ultimately ascertained at a wavelength of 570 nm, employing a microplate ELISA reader (Awareness Technology, Palm City, FL, USA).

### 2.6. Statistics

The experiments were conducted with a minimum of three repetitions. Numerical values are represented as the mean ± standard deviation (SD). Statistical analysis was executed on the exosomal miRNAs’ differential expression using the Student’s t-test via GraphPad Prism Version 9 (GraphPad Software, San Diego, Calif. USA). Statistical significance was set at p ≤ 0.05 and |log2FC| ≥ 1.0 were used to identify differentially expressed mRNAs between HC and RRMS patients.

## 3. Results

### 3.1. Exosome characterization

Serum and CSF samples of patients were obtained during the clinical meeting and subjected to pre-processing, as explicated in the methods section. Exosomes were obtained using the method of ultra-centrifugation and their visualization was carried out by means of transmission electron microscopy (TEM). TEM imaging has enabled the identification of a distinct population of nanovesicles with a range of 30-150 nm. It is noteworthy that these vesicles bear a cup-shaped morphology, which is characteristic of exosomes. The isolated exosomes were further examined through western blot analysis to identify exosomal markers. The extracted exosomes displayed a positive expression for CD9, CD81, and CD63 while exhibited a negative expression for calnexin, which is considered an exosome marker [32].

### 3.2. Exosomal miRNAs expression

A group of thirty individuals diagnosed with RRMS and an equivalent number of healthy control subjects were subjected to thorough investigation and analysis. The healthy group was carefully chosen based on the criteria of age and gender comparability with the RRMS patient group.

#### 3.2.1. Exosomal EBV-encoded miRNAs expression signatures in CSF

The relative amounts of ebv-miR-BART9-3p and ebv-miR-BART15 as sensitive biomarkers for EBV, not as disease-specific markers, were quantified using real-time polymerase chain reaction (qPCR). Interestingly, CSF exosomes from patients and control groups contained ebv-miR-BART9-3p and ebv-miR-BART15. However, the levels of both miRNAs were significantly increased in the CSF exosomes of patients with MS when compared to the control group (Fig. 1a and b). The presence of BART miRNAs in healthy individuals is in agreement with the abundant predisposition of healthy people to EBV infection, who have developed preventive measures to control and reduce the rate of infection.

**Fig. 1.**
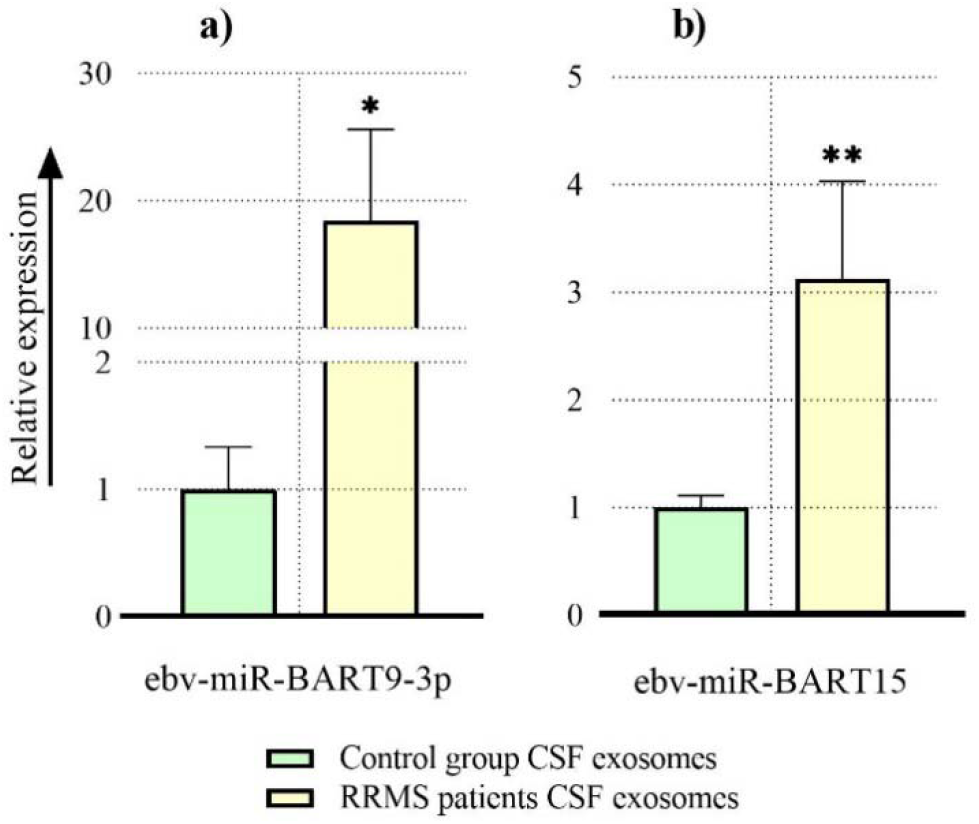
ebv-miRNAs expression in CSF exosomes of RRMS and control group. The expression level of ebv-miR-BART9-3p (a) and ebv-miR-BART15 (b) in CSF exosomes of RRMS patients were significantly higher than those of control group. The number of participants from each group was 30 individuals in the analyses (N=30). *: p<0.05; **: p<0.002.

#### 3.2.2. Expression of hsa-miR-21-5p and hsa-miR-146a-5p as EBV-stimulated human miRNAs

To further examine the association between EBV and MS, we studied the expression levels of two human miRNAs, hsa-miR-21-5p and hsa-miR-146a-5p, which are stimulated during EBV infection. Our results showed that the expression levels of hsa-miR-21-5p and hsa- miR-146a-5p were significantly upregulated in CSF samples compared to those in HC (Fig. 2a and b).

**Fig. 2.**
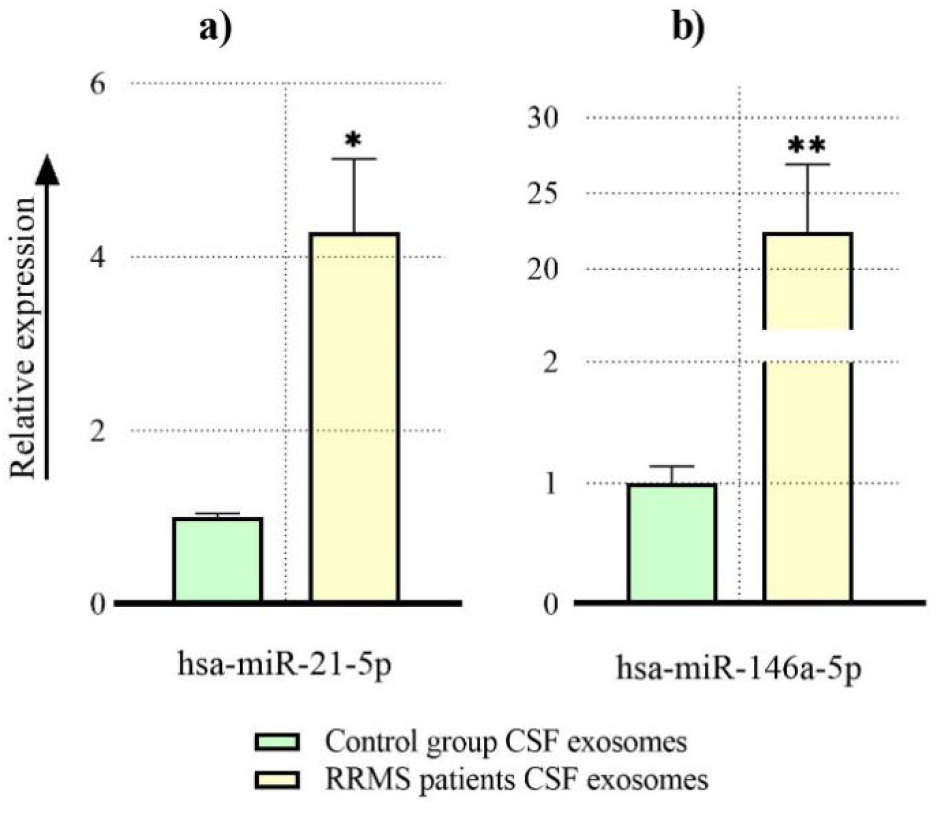
hsa-miR-21-5p and hsa-miR-146a-5p expression in CSF exosomes of RRMS and control group. The expression level of hsa-miR-21-5p (a) and hsa-miR-146a-5p (b) were remarkably increased in CSF exosomes of RRMS patients compared with control group. The number of participants from each group was 30 individuals in the analyses (N=30). *: p<0.05; **: p<0.002.

### 3.3. Exosomal inflammatory cytokines expression in CSF and serum

We also detected a significant increase in the levels of several inflammatory and autoimmune-related cytokines, including TNF-α, IL-23, IL-6, IL-1β, and IL-17, in CSF exosomes (Fig. 3) and those of serum (Fig. 4) in RRMS patients. Cytokines expression was more enhanced in CSF exosomes than in serum exosomes. The highest stimulation was observed in TGF-β (8.5-fold), followed by IL-23 (3.9-fold) in CSF exosomes.

**Fig. 3.**
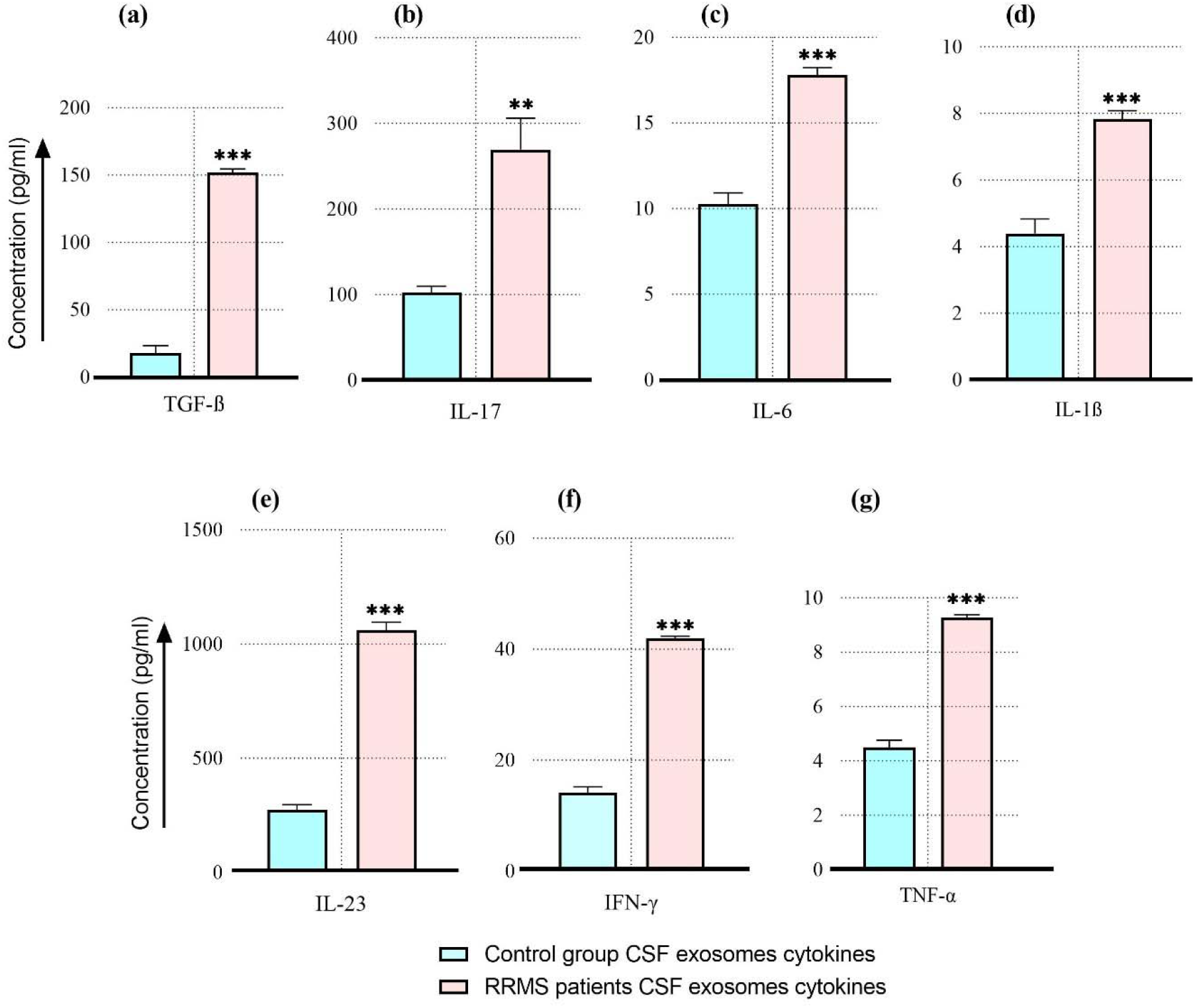
CSF exosomal cytokines profile of RRMS patients and control group. The number of participants from each group was 30 individuals in the analyses (N=30). *: p<0.05; **: p<0.002; ***: p<0.001.

**Fig. 4.**
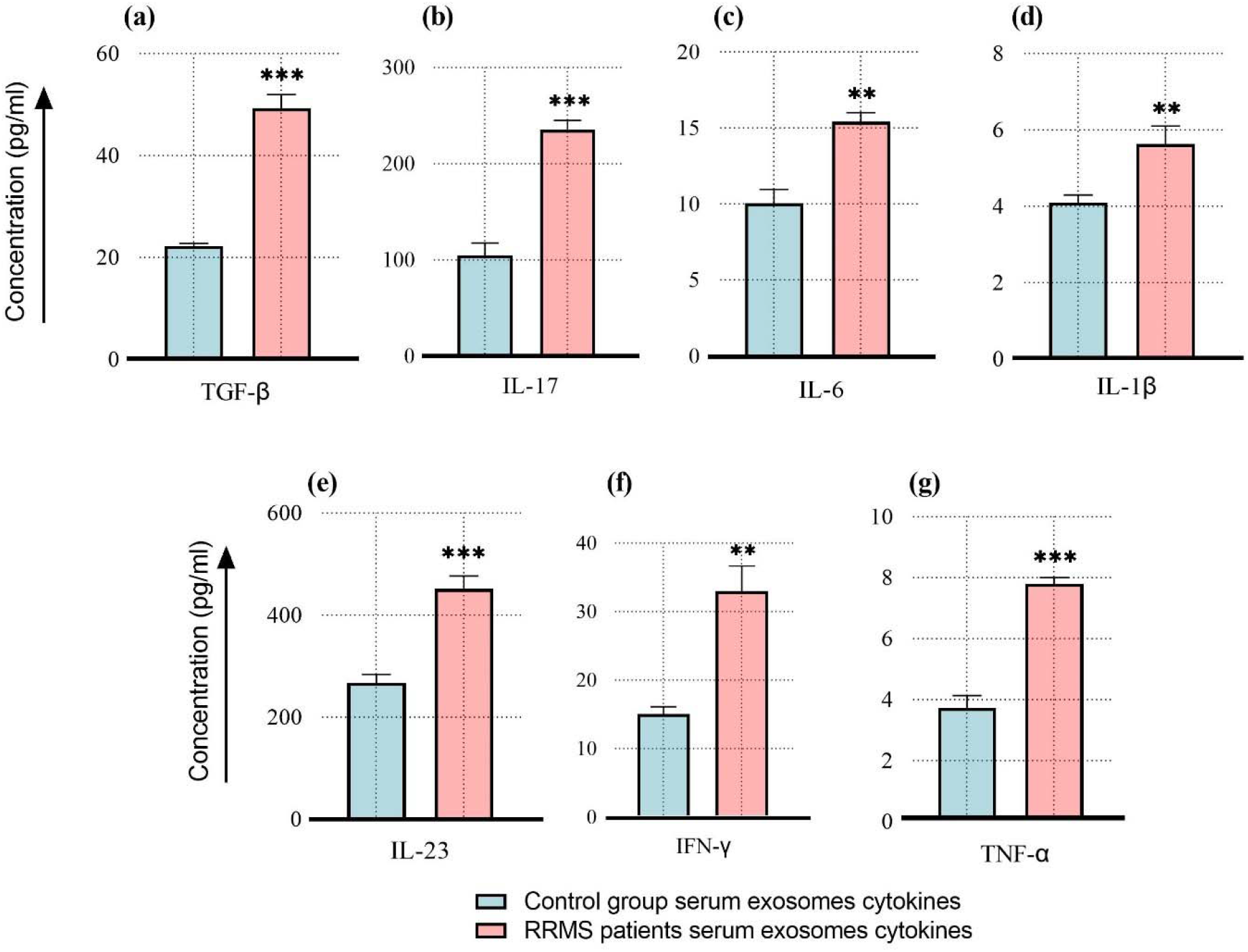
Serum exosomal cytokines profile of RRMS patients and control group. The number of participants from each group was 30 individuals in the analyses (N=30). *: p<0.05; **: p<0.002; ***: p<0.001.

## 4. Discussion

MS patients exhibit dysregulation of various cellular miRNAs. Disruption of miRNA homeostasis ultimately affects the levels of target mRNAs of genes involved in the local immune response. As a result, there is a tendency towards the inflammatory phenotype, which is evident in the elevated expression of IL17a, IFN-, and TNF-*α* mRNAs and their corresponding gene products [34]. Moreover, the increased level of MS-derived exosomal let-7i ultimately leads to the suppression of regulatory T cells, which are responsible for regulating the activity of pro-inflammatory T cell subsets [35]. Dysregulated miRNAs within cellular systems may serve as precursors for the infiltration of inflammatory cells and EBV-infected cells within the CNS. It is possible that miRNAs with EBV origin in the CNS of MS patients could also have an impact on the CNS resident cells, especially those involved in MS pathogenesis [36].

The observed significant increase in the levels of exosomal ebv-miR-BART9-3p (18.4- fold) and ebv-miR-BART15 (3.1-fold) in CSF implies a link between the elevation of EBV miRNAs and the triggering of MS disease. This finding is in line with Bjornevik and colleagues’ report [37], where they showed the positive correlation between EBV seropositivity before the onset of MS and documented EBV as the causal agent of MS. In other words, it seems that the increase in BARTs levels is the player in the infection-to-MS transition, not the presence of EBV miRNAs. Additionally, our findings support the significant role of EBV vaccination in the prevention of MS.

By taking into account that EBV infection is more prevalent in the CNS of RRMS patients compared to the control group [38], our data indicate that the site of EBV miRNAs elevation is the same as the site of pathogenesis and disease; CNS. Together, these results suggest greater demand for CNS-targeted therapies for RRMS patients. EBV-host interactions can result in dysregulation of host miRNA levels.

It has already been shown that EBV infection increases the expression of hsa-miR-21-5p and hsa-miR-146a-5p [39, 40]. Our results show that hsa-miR-21-5p and hsa-miR-146a-5p are upregulated in CSF exosomes of RRMS patients compared to HCs. This finding is by the upregulation of EBV-encoded BART9 and BART15 in CSF exosomes of RRMS patients compared to those of HCs.

It is shown that EBV directly influences cell signaling [41] and cytokine production, such as IL-1β, through the NF-κB pathway [27] in a dose-dependent manner [42]. EBV could induce secretion of IL-17, IL-23 [24], TNFα, IL-1β, IL-6, IFNγ [25–27], and TGF-β [28, 29]. Induction occurs in infected, and non-infected cells by direct interaction and indirectly via the supernatant and exosomes of the infected cells [25, 41]. Exosomes from monocyte-derived dendritic cells loaded with EBV peptides directly stimulate autologous CD8^+^ T cells to produce INF-β and TNF-α, even in the absence of dendritic cells [43]. EBV also enhances Th17 responses by stimulating pro-inflammatory cytokines such as IL-17 and IL-23 [24], which are involved in autoimmune diseases. Although several studies have evaluated cytokine expression in cell-free CSF and blood, none have reported the expression of cytokines in CSF and serum exosomes from patients with MS. Therefore, we aimed to study the expression levels of inflammatory cytokines in CSF and serum exosomes. Our results showed a significant increase in the levels of the studied cytokines in the exosomes. The observed elevation in the cytokine levels especially TGF-β (8.5-fold), TGF-β:TNF-α ratio from 4 to 16.4, and TGF-β:IFN-γ ratio from 1.3 to 3.6 in CSF exosomes of patients in comparison to the control group could be a result of EBV miRNA overexpression. Based on previous publications, all these cytokines exert pathogenic activity in MS [44]. TNF-α is an immunomodulatory cytokine, which is elevated in central and peripheral demyelinating lesions, serum, and brain plaques of MS patients [45, 46]. In line with these studies, we found a significant increase in the level of TNF-α in both CSF and serum of MS patients.

IL-23 is produced by activated dendritic cells and triggers the generation and survival of Th17 cells known to be crucial in chronic inflammation and autoimmune diseases. Th17 cells produce IL-17, which has been detected in MS lesions. IL-23, IL17, and Th17 cells, have all been implicated in the pathogenesis of MS [47]. The local synthesis of IL-23 induces a proinflammatory milieu with upregulation of cytokines and several complement factors, leading to demyelination [48]. Studies have suggested that IFN-γ exerts pathogenic effects in MS [44]. IFN-γ is largely produced by CD8+ cytotoxic T cells, which have been reported to be present and to have a pathogenic profile in the MS CNS [49]. Previous studies have shown the presence of T lymphocytes that produce IFN-γ in the perivascular infiltrates of patients with MS [49, 50]. Other sources of IFN-γ are Th1, Th17, and NK cells, all of which play important roles in the pathogenesis of MS [44, 51, 52]. TGF-β is expressed in CNS and can alter the phenotype or function of effector T cells that infiltrate the CNS in the context of CNS infections and autoimmunity [53]. The significance of TGF-β in MS has been highlighted by the observations that TGF-β promotes the differentiation of murine Th17 cells in the presence of IL-6 and IL-23 [54, 55]. Interestingly, our findings show the upregulation of IL-6 as well as IL-17 in both CSF and serum of MS patients. Based on these results, it is concluded that there is a crosstalk between the studied BARTs elevation and inflammatory cytokines upregulation in RRMS patients. This explains the causal role of EBV in disease onset and the infection-to-MS transition.

## Funding

This study was supported by the Molecular Medicine Research Center, Tabriz University of Medical Sciences, (number 62877), Iran.

## CRediT authorship contribution statement

Conceptualization and supervision: M.S.H. Patients’ selection: H.A., M.T.

Performing experiments: M.M., S.M.

Data validation and statistical analysis: M.M., S.M., V.H., H.K., O.M., H.A., M.S.H. Writing: M.M., S.M., M.S.H.

## Ethical declaration

This study was performed in line with the principles of the Declaration of Helsinki. Approval was granted by the Ethics Committee of Tabriz University of Medical Sciences (IR.TBZMED.REC.1402.348).

## Informed consent

Informed consent was obtained from all individual participants included in the study.

## Declaration of competing interest

The authors declare that they have no conflict of interest.

## Acknowledgment

The authors greatly appreciate Professor Lawrence Steinman’s support, constructive input, and assistance in editing the manuscript. This is a report of the database from a Ph.D. thesis registered in Tabriz University of Medical Sciences with the Number 62877.

## Data availability

All data are available in the main text.

## Consent to publish

The authors affirm that human research participants provided informed consent for publication.

## Notes

### Competing Interest Statement

The authors have declared no competing interest.

