## Supplemental File for "Epstein-Barr virus-encoded BART9 and BART15 miRNAs are elevated in exosomes of cerebrospinal fluid from relapsing-remitting multiple sclerosis patients"

**Supplementary material**

**Materials and methods**

**Size and morphology of exosome**

The isolated exosomes were visualized by transmission electron microscopy (TEM) to evaluate the shape and size of the exosomes. three-hundred mesh carbon-coated copper grids were incubated for 5 minutes. All copper grids were negatively stained with 1.5% (w/v) uranyl acetate for 2 minutes and left overnight. The copper grids were observed and photographed using Zeiss LEO 906 instrument (Freiburg, Switzerland) with an acceleration voltage of 80 kV**.**

**Western blotting**

Equal amounts of exosomes were subjected to lysis in protein lysis buffer and western blot analyses followed according to the method of Mohammadinasr et al. [1]. The identity of the exosomes was confirmed through the detection of exosomal markers: anti-CD9 antibody (sc-13118, Dallas, USA), anti-CD63 antibody (sc-5275, Dallas, USA), and anti-CD81 antibody (sc-166029, Dallas, USA). In addition, mouse monoclonal anti-calnexin (sc-23954, Dallas, USA) was used as a negative control.

**RNA isolation**

Exosomal total RNA was isolated using the TRIzol method (RiboEx.LS, Seoul, Korea). In order to lyse the samples, 750 µl of RiboEx and 200 µl of chloroform were added and the resultant mixture was incubated at room temperature for two minutes. Phase separation was done by centrifuging samples at 12,000 x g for 20 minutes at 4˚C. The upper aqueous phase was collected for RNA extraction. One volume of isopropanol was added to the aqueous phase. Samples were incubated at -20˚C for 90 minutes to facilitate RNA precipitation. Post incubation, samples were centrifuged at 12,000 x g for an hour at 4˚C to pellet the RNA. Subsequently, RNA pellets were washed with one ml of 75% ethanol and centrifuged at 12,000 x g for 20 minutes at 4˚C. The pellet was air-dried for 10 min and re-suspended in 20 μL of DEPC-treated water (Cinnagen, Tehran, Iran). RNA concentration and quality were determined using a Nanodrop Spectrophotometer (Thermo Scientific NanoDrop, USA).

**Results**

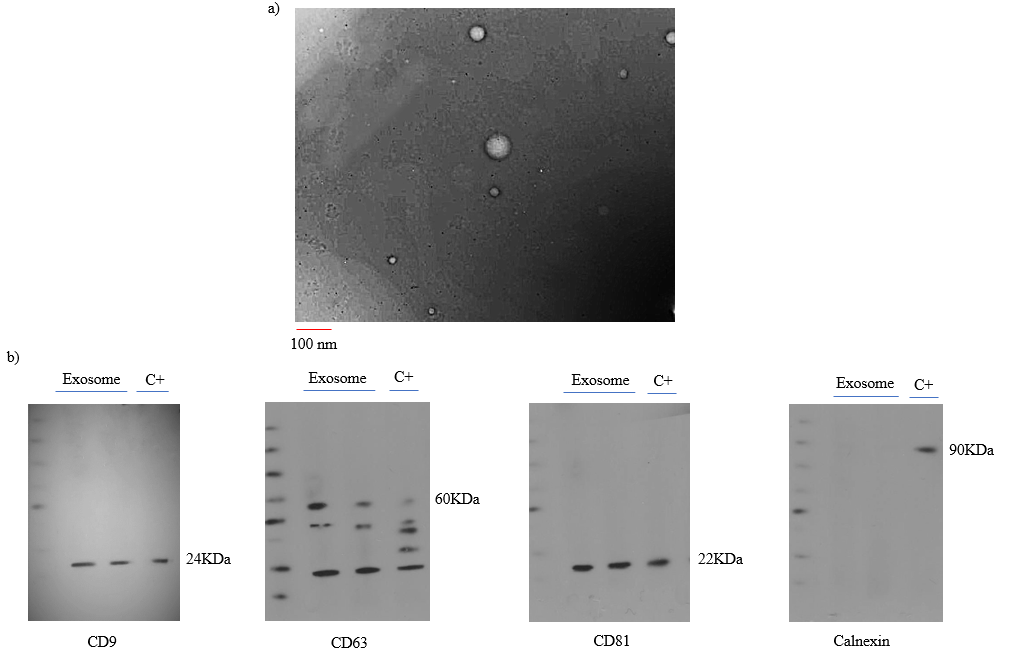

**Fig. 1. The detection and characterization of exosomes in serum and CSF.** (a) The morphology and size of exosomes in the cerebrospinal fluid (CSF) were studied using transmission electron microscopy (TEM). The results showed that the exosomes have cup-shaped bodies with a diameter of 50 to 150 nm. (b) Exosomes in blood and CSF were identified using specific protein markers. Serum and CSF exosomes were positive for CD9, CD63, and CD81 markers and negative for calnexin. Cell lysate (C+) used as positive control.

**Table 1. Patients and healthy individuals’ demographic characteristics in this study.**

| **Demographics information** | **Patients** | **HC (Serum)** | **HC (CSF)** |
| --- | --- | --- | --- |
| Male/Female | 9/21 | 11/19 | 11/19 |
| Age (± SD) | 43.0 (12.9) | 40.3 (11.8) | 40.6 (11.4) |
| EDSS (± SD) | 1.5 (1) |  |  |
| Oligoclonal Band Numbers | 6.26 (3.19) |  |  |
| Lesion (identified by MRI) |  |  |  |
| Treatment |  |  |  |

Abbreviations: HC, Healthy Control; CSF, Cerebrospinal Fluid; SD, Standard Deviation; EDSS, Expanded Disability Status Score
